# Recruitment of PI4KIIIβ to the Golgi by ACBD3 is dependent on an upstream pathway of a SNARE complex and golgins

**DOI:** 10.1101/2023.04.13.536018

**Authors:** Danièle Stalder, Igor Yakunin, David C. Gershlick

## Abstract

ACBD3 is a protein localised to the Golgi apparatus and recruits other proteins, such as PI4KIIIβ, to the Golgi. However, the mechanism through which ACBD3 itself is recruited to the Golgi is poorly understood. This study demonstrates there are two mechanisms for ACBD3 recruitment to the Golgi. First, we identified that an MWT^374-376^ motif in the unique region upstream of the GOLD domain in ACBD3 is essential for Golgi localisation. Second, we use unbiased proteomics to demonstrate that ACBD3 interacts with SCFD1, a Sec1/Munc-18 (SM) protein, and a SNARE protein, SEC22B. CRISPR-KO of SCFD1 causes ACBD3 to become cytosolic. We also found that ACBD3 is redundantly recruited to the Golgi apparatus by two golgins: golgin-45 and giantin, which bind to ACBD3 through interaction with the MWT^374-376^ motif. Taken together, our results demonstrate that ACBD3 is recruited to the Golgi in a two-step sequential process, with the SCFD1-mediated interaction occurring upstream of the interaction with the golgins.

## Introduction

The Golgi apparatus is a eukaryotic evolutionarily conserved organelle with a central role in trafficking, processing, and sorting newly synthesised proteins and lipids^1–3^. The metazoan Golgi apparatus is a collection of Golgi stacks, with 10-20 per cell. Each Golgi stack consists of 5-7 connected cisternae^4^. The cisternae are polarised such that there is a *cis* and *trans* cisterna. *De novo* biosynthetic protein products are delivered to the *cis*-face of the Golgi apparatus and exit at the *trans*-face or the *trans*-Golgi network (TGN). Within the Golgi apparatus, proteins are sorted into export domains for recycling back to the endoplasmic reticulum, delivery to the plasma membrane or delivery to the endolysosomal system. Accordingly, the Golgi apparatus represents a central sorting hub for protein trafficking and is essential for the sorting and delivering of many important proteins. Structural and functional changes to the Golgi apparatus are linked to numerous diseases, including developmental defects, neurodegenerative and infectious diseases, and certain cancers^3,5^.

Phosphoinositides are a class of phospholipids embedded in lipid bilayers in eukaryotic membranes^6^. Phosphatidylinositol, the precursor to all phosphoinositides, consists of two fatty acid chains attached to a glycerol backbone and an inositol head group. The inositol head group can be phosphorylated at the free hydroxyl groups at positions D3, D4, and D5 to form a large class of various phosphoinositides. These distinct phosphoinositides have a role in compartment identity and have different localisations within the endomembrane system. PI4P is crucial in the Golgi apparatus for driving several key processes through recruiting PI4P-interacting proteins^7-9^. Examples of PI4P recruited proteins include GOLPH3^10,11^ and arfaptins^12^, which allow intra-Golgi trafficking, and clathrin adaptor proteins at the *trans*-Golgi, such as the GGA family^13^ and AP1^14^. Lipid transport proteins, such as OSBP, are also partly recruited by PI4P^15-18^. Although the most characterised role of PI4P on the Golgi apparatus is at the TGN and *trans*-Golgi, PI4P is most enriched on the *cis*-Golgi apparatus and is demonstrated to have a central role in the delivery of COPII vesicles from the ER to the *cis*-Golgi^19,20^.

PI4KIIIβ has the best understood role in generating PI4P at the Golgi apparatus^7^. PI4KIIIβ phosphorylates phosphatidylinositol at the D4 position. One way PI4KIIIβ is recruited to the Golgi apparatus is by Acyl-CoA Binding Domain Containing 3 (ACBD3, also known as GCP60 and PAP7)^21^. Loss of ACBD3 is lethal in mice, and ACBD3 is upregulated in multiple breast cancers in which high ACBD3 expression correlates with poor patient survival^22,23^. ACBD3 is a cytosolic protein mainly localised to the Golgi apparatus and is important for Golgi organisation^24,25^. ACBD3 consists of an ACBD domain, a CAR-Q domain, a ‘unique region’, and a GOLD domain. ACBD3 is considered a multi-functional scaffolding protein and interacts with various proteins^26,27^, including PPM1L^28^, and TBC1D22,a Rab33b GTPase activating protein (GAP)^29^. The ACBD domain oligomerises upon binding to C18:1-CoA or C16:0-CoA^30^ and recruits the membrane-shaping protein FAPP2^25^, the CAR-Q domain recruits PI4KIIIβ^31^, and the GOLD domain and its extended unique reigon interact with multiple different golgins^29,32-34^ (giantin, golgin-45 and golgin-160). The recruitment of ACBD3 to the Golgi apparatus, therefore, represents a crucial functional step in the homeostasis and regulation of the Golgi apparatus.

There are multiple proposed mechanisms for the recruitment of ACBD3 to the Golgi apparatus. Initially, based on a yeast-2 hybrid interaction, the golgin giantin was proposed to recruit ACBD3^32^ ; however, the knock-down of giantin does not result in the loss of ACBD3 recruitment to the Golgi apparatus^29^. Recently, the loss of ARL5B was demonstrated to cause the loss of the Golgi pool of ACBD3^35^ ; however, ARL5B is *trans*-Golgi localised compared to the *cis*- and *trans*-Golgi localised ACBD3. Therefore, the mechanism by which ACBD3 is recruited to the Golgi apparatus remains unclear.

Here, we show that ACBD3 is recruited to the Golgi apparatus by an MWT^374-376^ motif in the ‘unique region’ upstream of the GOLD domain. We identify a host of new Golgi localised ACBD3 interactors using unbiased proteomics. We demonstrate that the SNARE SEC22B and associated SM protein SCFD1 interact with ACBD3 and abrogation of SCFD1 causes loss of ACBD3 from the Golgi apparatus. We also identify that the golgins giantin and golgin-45 interact with ACBD3 dependent on the MWT^374-376^ motif, and loss of both golgins results in loss of ACBD3 localisation, indicating a redundant function. Our data support a novel two-step mechanism for ACBD3 recruitment to the *cis*-Golgi apparatus.

## Results

### ACBD3 is recruited to the Golgi apparatus via a protein-protein interaction within a unique region of the GOLD domain

ACBD3 is a 528 amino acid protein containing ACBD, CAR-Q, and GOLD domains (Figure 1A) that has been reported to localise to the Golgi apparatus^24,25,32,36^. We performed high-resolution Airyscan imaging of HeLa cells immunolabeled for ACBD3 and *cis* and *trans*-Golgi markers. Despite having no transmembrane domain or other known Golgi interacting domain^32^, ACBD3 showed significant colocalisation with *cis* and *trans*-Golgi markers (Figure S1A and B), consistent with previous findings^35,36^.

**Figure 1.**
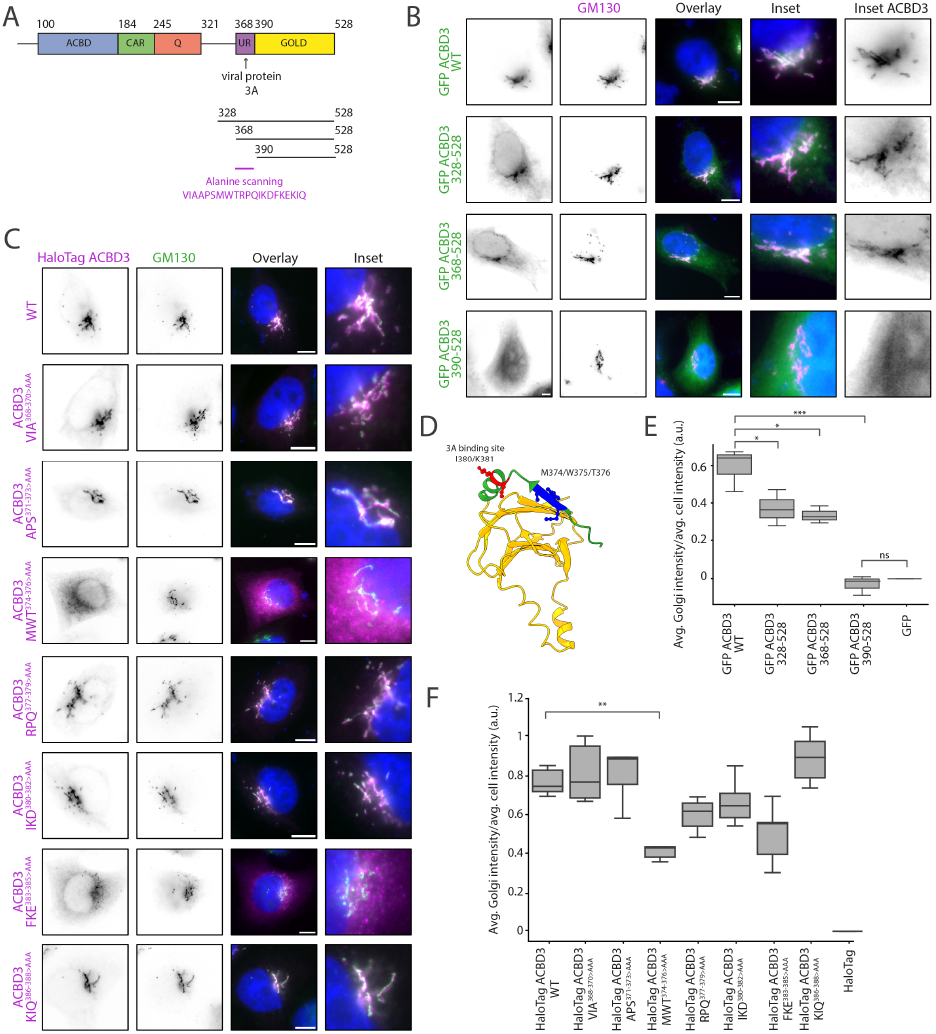
The unique region of ACBD3 is essential to target ACBD3 to the Golgi apparatus. (**A**) Schematic of the domain organisation of ACBD3. From N- to C-terminus, ACBD3 has a acyl-CoA binding domain (ACBD), a charged amino acid region (CAR) domain, a glutamine-rich domain (Q-domain), a unique region (UR), and a Golgi dynamic domain (GOLD domain). To investigate the importance of the UR for the Golgi localisation of ACBD3, we generated different truncations of the UR-GOLD domain of ACBD3 and the subsequent alanine scanning approach in the UR is indicated (**B-C**) The UR is essential to target ACBD3 to the Golgi apparatus. Widefield imaging of HeLa cells expressing different truncations of GFP-ACBD3 (B) or different HaloTag-ACBD3 constructs mutated in the UR (alanine scanning mutagenesis) (C). The Golgi was stained with GM130 antibody and the nucleus with DAPI. Scale bar: 10 μm. (**D**) Structure of the GOLD domain (yellow) and the unique region (green) with MWT (M374/W375/T376) residues highlighted in blue and protein 3A targeted residues^39^ in red (I380/K381) (AlphaFold2). (**E and F**) Quantitative analysis with a CellInsight CX7 high-content microscope of the Golgi localisation of the different truncations and mutants of ACBD3 (B and C). The ratio between the average intensity of ACBD3 at the Golgi and in the total cell normalised to GFP or HaloTag alone is indicated (a.u). The experiments were performed 3 to 5 times independently. Tukey’s multiple comparisons test (HSD, FWER=0.05) was performed. *P≤0.05; **P≤0.01; ***P≤0.001; ns: not significant.

To identify the domain by which ACBD3 is recruited to the Golgi apparatus, we examined other known ACBD3 recruitment factors. Picornavirus 3A peptide recruits ACBD3 to viral replication sites through a protein-protein interaction via the ‘unique region’ (UR) of the GOLD domain of ACBD3^21,37-44^ (Figure 1D). We reasoned that the 3A peptide must outcompete the endogenous Golgi-localised ACBD3 recruitment factor. To test this hypothesis, we generated truncations of ACBD3 in this region (Figure 1A). GFP-fused truncations that included the GOLD domain and the UR (328-528, and 368-528) are still localised to the Golgi apparatus (Figure 1B). However, the isolated GOLD domain, without the ‘unique region’, is mainly cytosolic. To confirm our findings quantitatively, we built an unbiased high-throughput imaging pipeline to assess Golgi localisation (Figure 1E). The different ACBD3 GFP-fused truncations were assessed for colocalisation with Golgi marker GM130. Our quantitative approach confirms our observation that the UR upstream of the GOLD domain is essential for the Golgi localisation of ACBD3.

We performed alanine scanning mutagenesis of the 21 amino acids of the UR (Figures 1A, 1C, and 1D) and assessed their Golgi localisation as before (Figures 1C,1F). Most of the mutants demonstrated a localisation comparable to WT ACBD3 aside from one mutant, MWT^374-376>AAA^ (Figure 1D), which was relocalised to the cytosol. We thus demonstrated that the residues MWT^374-376^ in the UR of the GOLD domain of ACBD3 participate in a protein-protein interaction that recruits ACBD3 to the Golgi apparatus.

### ACBD3 interacts with an array of Golgi localised proteins and knock-out of SCFD1 causes ACBD3 to be mislocalised to the cytosol

To identify the factors responsible for the recruitment of ACBD3 to the Golgi apparatus, we performed immunoisolation of GFP-ACBD3 or the negative control GFP using GFP-nanobody conjugated agarose beads and assessed using mass spectrometry.

By definition, the unknown protein responsible for ACBD3 recruitment to the Golgi apparatus localises to the Golgi apparatus. Therefore, interactors were bioinformatically filtered for Golgi localisation using publicly available datasets (Figure 2A). Identified putative interactors included previously identified ACBD3 interactors giantin (GOLGB1) and golgin-45 (BLZF1).

**Figure 2.**
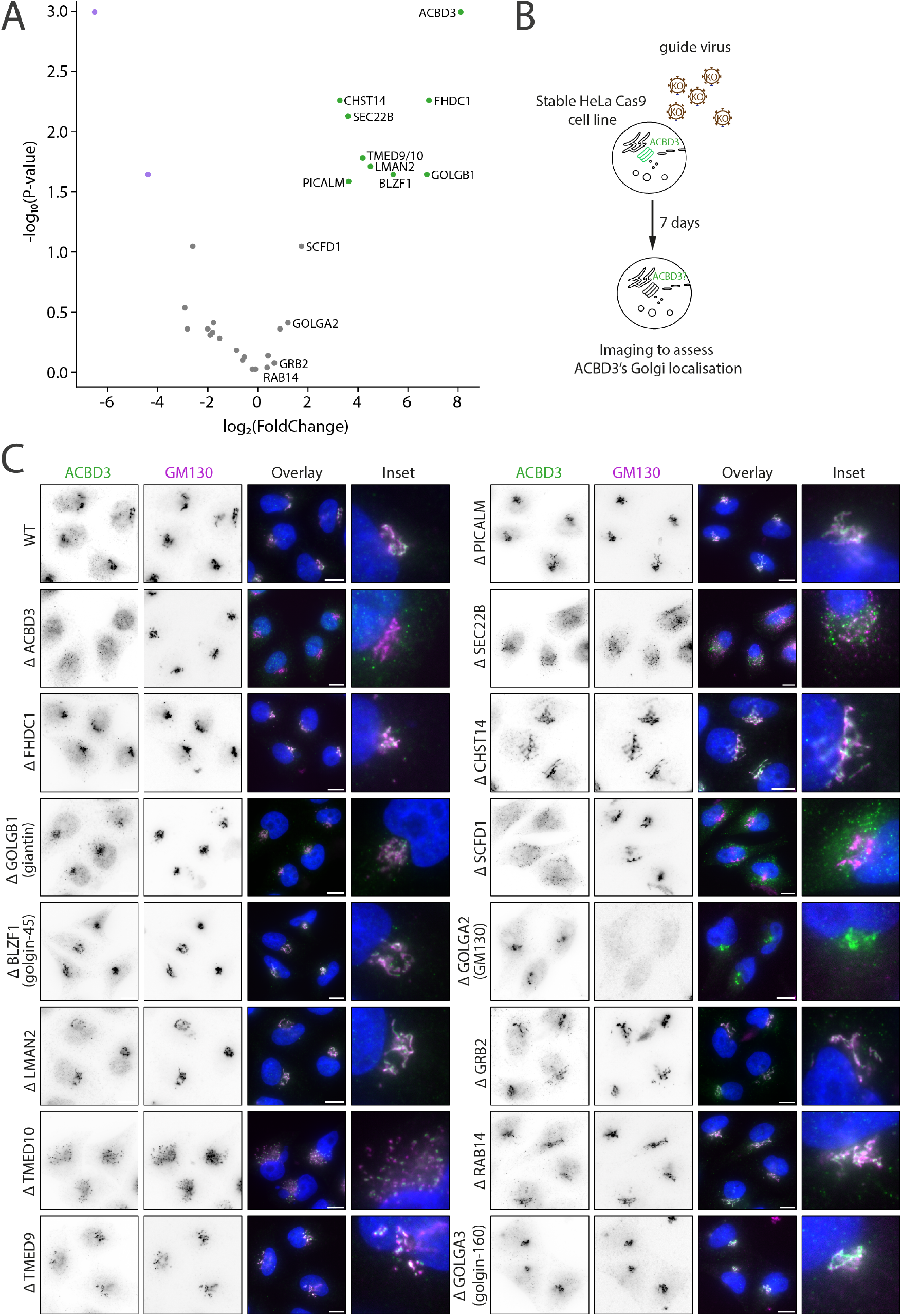
Identification of new interactors of ACBD3. (**A**) Volcano plot illustrating Golgi proteins identified as potential binding partners of ACBD3. Three independent GFP-trap immunoprecipitation experiments were performed on HEK-293T cells overexpressing GFP-ACBD3 or GFP alone as a control. The resultant immunoprecipitates were analysed by mass spectrometry, and Golgi-associated proteins were identified. (**B**) Schematic representation of the follow-up transient CRISPR/Cas9 KO screen with the 13 first hits of proteins identified by mass spectrometry. Golgin-160 (GOLGA3), an interactor of ACBD3, was included in the screen. HeLa cells were infected with the according guides and imaged 7 days post-infection. (**C**) Widefield imaging of endogenous ACBD3 in the different HeLa KO cells. Endogenous GM130 is stained to verify the Golgi integrity. Scale bar: 10 μm. Nucleus stain = DAPI.

To determine if these putative interactors were necessary for the ACBD3 recruitment to the Golgi apparatus, ‘hits’ were individually knocked-out (KO) using CRISPR-Cas9 in HeLa cells (Figure 2B and C). The depletion of target proteins was verified by qPCR analysis (Figure S2A). As previously shown, in WT cells, ACBD3 colocalised with GM130 (Figure 2C). In the positive control, CRISPR-Cas9 guides targeting ACBD3 caused loss of ACBD3 staining at the Golgi apparatus (Figure 2C and quantified in S2B). Most of the KOs showed no observable effect, and the loss of giantin did not affect the recruitment of ACBD3 to the Golgi. Loss of TMED10 and SEC22B caused a drastic loss of Golgi organisation resulting in fragmented Golgi apparati. Loss of SCFD1, however, resulted in the almost complete loss of ACBD3 from the Golgi apparatus (Figure 2C and quantified in S2B). Therefore, we conclude that SCFD1 is a previously unidentified essential factor for the recruitment of ACBD3 to the Golgi apparatus.

### SCFD1 and ACBD3 interact upstream of PI4KIIIβ recruitment to the Golgi apparatus

SCFD1 (aka SLY1) is a soluble cytosolic Sec1/Munc18-like protein recruited to the *cis*-Golgi apparatus to participate in catalysing the formation of a SNARE complex^45-47^. Exogenous overexpression of an SCFD1-HaloTag fusion colocalised with endogenous ACBD3 at the *cis*-Golgi apparatus (Figure 3A).

**Figure 3.**
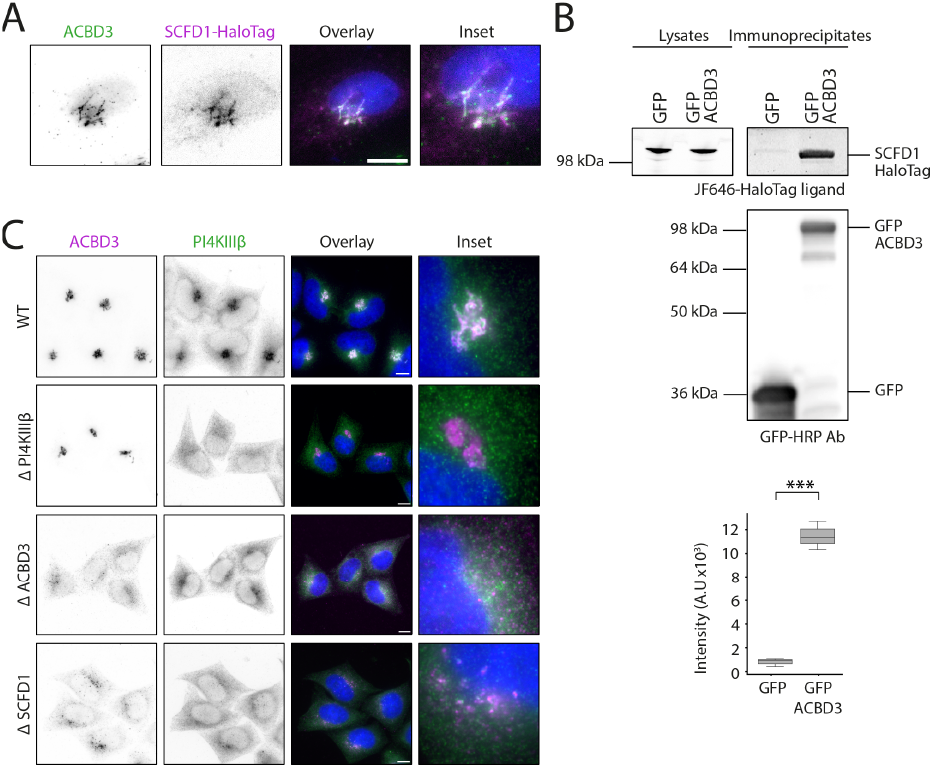
ACBD3 interacts and colocalises with SCFD1. **(A)** SCFD1 colocalises with ACBD3 at the Golgi. Widefield imaging of HeLa cells transfected with SCFD1-HaloTag and stained for endogenous ACBD3. Nucleus stain = DAPI. Scale bar: 10 μm. (**B**) ACBD3 interacts with SCFD1. GFP alone or GFP-ACBD3 was immunoprecipitated with GFP-nanobody conjugated agarose beads from HEK-293T cells co-expressing SCFD1-HaloTag. Bound SCFD1-HaloTag and the amount of SCFD1-HaloTag in 0.125% of the lysates were detected by direct imaging of the JF646 HaloTag ligand dye on the gel. The amount of GFP and GFP-ACBD3 in the immunoprecipitates was detected with an anti-GFP HRP conjugate antibody. The experiment was repeated three times, and the intensity of the bands was quantified using Fiji. The mean and SD of three independent experiments are shown and a two-tailed t-test was performed on the data. ***P ≤ 0.001. (**C**) SCFD1 is important for the recruitment of PI4KIIIβ to the Golgi apparatus. Widefield imaging of endogenous ACBD3 and PI4KIIIβ in CRISPR-Cas9 KO cells of PI4KIIIβ, ACBD3 and SCFD1. Nucleus stain = DAPI. Scale bar: 10 μm.

Co-expression of SCFD1-HaloTag with GFP-ACBD3, or the negative control GFP and subsequent immunoprecipitation with GFP-nanobody conjugated agarose beads corroborated mass spectrometry data that SCFD1 interacts with ACBD3 (Figure 3B).

ACBD3 recruits and activates PI4KIIIβ at the Golgi apparatus^21,31^. Using immunostaining against endogenous proteins, we confirmed that ACBD3 colocalises with PI4KIIIβ (Figure 3C). CRISPR-KO of PI4KIIIβ did not affect the localisation of ACBD3 (Figure 3C). In agreement with previous literature, the abrogation of ACBD3 caused a loss of localisation of PI4KIIIβ to the Golgi apparatus^21^ (Figure 3C). CRISPR-Cas9 KO of SCFD1 resulted in the loss of both ACBD3 and PI4KIIIβ at the Golgi apparatus. Thus, SCFD1 is upstream of ACBD3, which is upstream of PI4KIIIβ.

It has been proposed that SCFD1 interacts with the *cis*-Golgi SNARE STX5 allowing it to switch to an open conformation by exposing its SNARE domain^46,47^. Then, SCFD1 bridges STX5 to the incoming COPII vesicles and promotes the formation of the SNARE complex between STX5 and the v-SNARE SEC22B, ultimately leading to membrane fusion. Interestingly, we also identified SEC22B as a binding partner of ACBD3 in our mass spectrometry experiment (Figure 2A).

### The GOLD domain of ACBD3 interacts with the longin domain of SEC22B

To confirm the interaction of ACBD3 with SEC22B, we performed co-immunoprecipitation experiments in cells expressing GFP or GFP-ACBD3 in combination with HaloTag-SEC22B (Figure 4A). GFP immunoisolation validated our mass spectrometry data that ACBD3 interacts with SEC22B.

**Figure 4.**
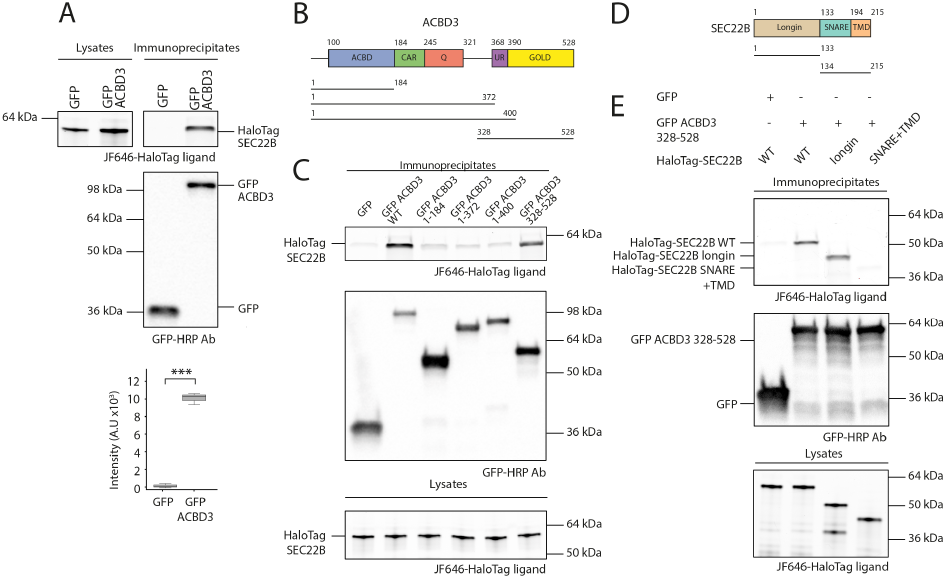
The GOLD domain of ACBD3 interacts with the longin domain of SEC22B. (**A**) ACBD3 interacts specifically with SEC22B. GFP alone or GFP-ACBD3 was immunoprecipitated with GFP-nanobody conjugated agarose beads from HEK-293T cells co-expressing HaloTag-SEC22B. Bound HaloTag-SEC22B and the amount of HaloTag-SEC22B in 0.125% of the lysates were detected by directly imaging the JF646 HaloTag ligand on the gel. The amount of GFP and GFP-ACBD3 in the immunoprecipitates was detected with an anti-GFP HRP conjugate antibody. The experiment was repeated three times and the intensity of the bands was quantified using Fiji. The mean and SD of three independent experiments are shown and a two-tailed t-test was performed on the data ***P≤0.001. (**B and D**) Schematic of the different truncations of ACBD3 and SEC22B. (**C**) The GOLD domain of ACBD3 interacts with SEC22B. Different GFP-ACBD3 truncations were co-expressed with HaloTag-SEC22B in HEK-293T cells and immunoprecipitation experiments were performed as described in A. (**D**) The GOLD domain of ACBD3 interacts with the longin domain of SEC22B. GFP and GFP-ACBD3 328-528 were co-expressed with different truncations of HaloTag-SEC22B in HEK-293T cells and immunoprecipitation experiments were performed as described in A.

To identify which domain of ACBD3 interacts with SEC22B, we made a series of truncations to GFP-ACBD3 and performed co-immunoprecipitation experiments (Figure 4B-C). We have therefore identified that ACBD3 interacts with SEC22B by the UR and GOLD domain (328-528).

SEC22B contains transmembrane, SNARE, and longin domains (Figure 4D). The transmembrane domain is buried in the lipid bilayer precluding interaction with ACBD3. To know if ACBD3 interacts with the SNARE or longin domains, HaloTag-SEC22B truncations were generated, and GFP-nanobody immunoisolation was performed as before (Figure 4E). As shown above, full-length SEC22B interacts with GFP-ACBD3 (328-528). The SNARE and transmembrane domain did not interact with ACBD3 (328-528); however, the longin domain alone does interact with ACBD3 (328-528). We thus conclude that the UR and GOLD domain of ACBD3 interacts with the longin domain of SEC22B and that ACBD3 is part of the SNARE complex involving the SM protein SCFD1 and the v-SNARE SEC22B.

### The UR in ACBD3 does not mediate the interaction to SCFD1 or SEC22B

These results raised the possibility that the MWT^374-376^ residues in the UR of ACBD3, interact with SCFD1 and SEC22B to recruit ACBD3 to the Golgi apparatus. We, therefore, repeated the GFP-nanobody immunoprecipitation with a series of truncations in the UR-GOLD domain of ACBD3 used previously (Figure 1A), and we also included the ACBD3 MWT^374-376>AAA^ mutant, which results in the mislocalisation of ACBD3 (Figure 5A-B). Surprisingly, SCFD1 and SEC22B interact strongest with the cytosolic truncation of GFP-ACBD3, which does not contain the UR (390-528) of the GOLD domain, and with the MWT^374-376>AAA^ mutant. Therefore, there must be a second mechanism by which ACBD3 is recruited to the Golgi apparatus.

**Figure 5.**
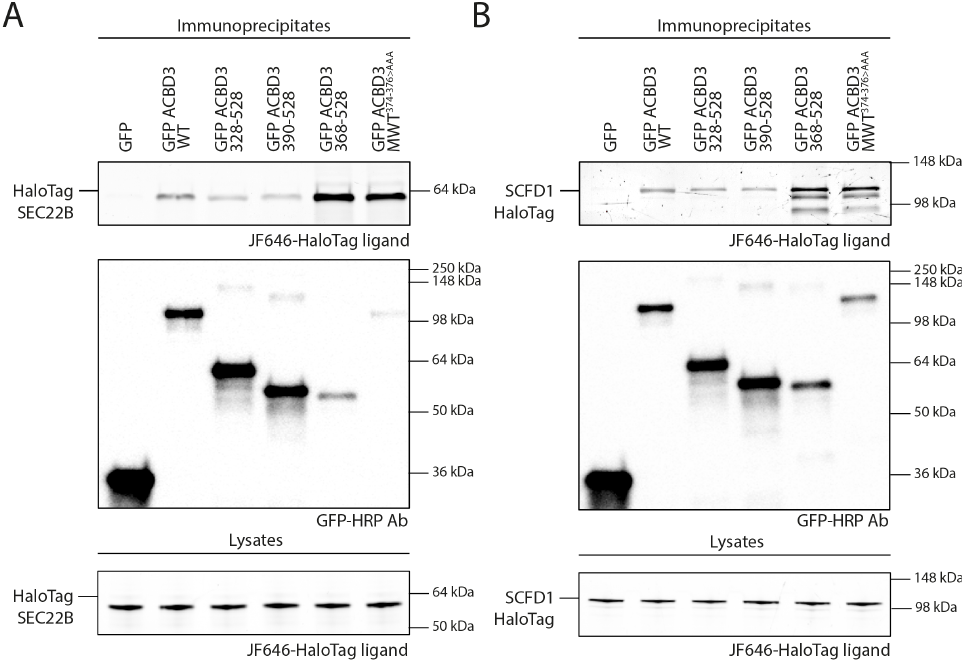
SEC22B and SCFD1 do not interact with MWT^374-376^ residues in the unique region. (**A and B**) Different truncations of GFP-ACBD3-UR-GOLD domain and GFP-ACBD3 MWT^374-376>AAA^ mutant were co-expressed either with HaloTag-SEC22B (A) or SCFD1-HaloTag (B) in HEK-293T cells and immunoprecipitation experiments were performed as described above.

### Golgin-45 and giantin redundantly recruit ACBD3 to the Golgi apparatus

To understand the additional mechanism by which ACBD3 is recruited to the Golgi apparatus, we reassessed the Golgi interactome of ACBD3. Giantin (GOLGB1) has been previously demonstrated to interact with ACBD3, and we also identified it in our interactome. We also observed an interaction with golgin-45 (BLZF1), which has also been previously documented^29^. Finally, other reports show that golgin-160 interacts with ACBD3^33,34^, although we did not observe this in our dataset. Golgins have been documented to act redundantly^48^, so we hypothesised that, in addition to SCFD1, ACBD3 was being recruited to the Golgi apparatus by a combination of golgins.

To test if combinations of golgins could be redundantly recruiting ACBD3 to the Golgi apparatus, we used CRISPR-Cas9 to KO combinations of the three golgins (Figure 6A). KO of giantin and golgin-160, or golgin-45 with golgin-160, did not cause a loss of ACBD3 at the Golgi. However, KO of both golgin-45 and giantin drastically affects the localisation of endogenous ACBD3 at the Golgi apparatus (Figure 6A and quantified in S2B-C). This indicates that the second recruitment mechanism of ACBD3 to the Golgi apparatus is via two redundant golgins, golgin-45 and giantin.

**Figure 6.**
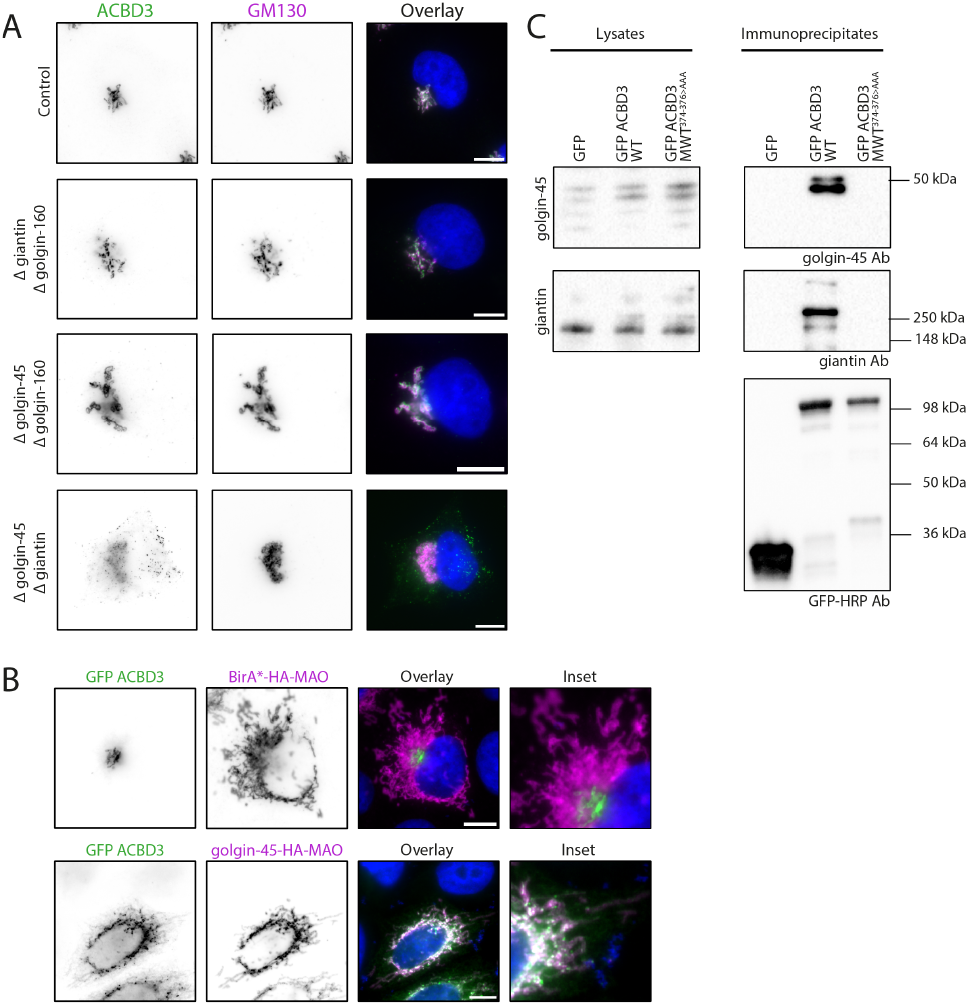
Golgin-45 and giantin interact with ACBD3 through MWT^374-376^ residues and are essential for its Golgi localisation. (**A**) The double KO of golgin-45 and giantin affects ACBD3 localisation. Wide-field imaging of endogenous ACBD3 and GM130 in CRISPR-Cas9 KO cells of golgins. Scale bar: 10 μm. Nucleus stain = DAPI. (**B**) Golgin-45 is sufficient to recruit ACBD3 to the mitochondria. Mitotrap assay with co-expression of GFP-ACBD3 with golgin-45-HA-MAO. BirA*-HA-MAO was used as a negative control. Scale bar: 10 μm. Nucleus stain = DAPI. (**C**) MWT^374-376^ residues are essential for the interaction of ACBD3 with golgins. GFP alone, GFP-ACBD3 or GFP-ACBD3 MWT^374-376>AAA^ were overexpressed in HEK-293T cells. Immunoprecipitation experiments were performed and immunoprecipitates were blotted for endogenous giantin and golgin-45.

To investigate if there is true functional redundancy between both golgins we sought to test if golgin-45 alone is sufficient for the recruitment of ACBD3 to intracellular organelles. To do so, we used the mitotrap assay in which golgins are ectopically localised to the mitochondrial outer membrane, where they can still capture vesicles^49^ (Figure 6B). Negative control of an exogenous enzyme BirA* targeted to the mitochondria did not redistribute GFP-ACBD3 to the mitochondria. Ectopic expression of golgin-45 localised to the mitochondria caused a subset of the GFP-ACBD3 to be relocalised to the mitochondria from the Golgi apparatus. These data indicate that golgin-45 alone is sufficient to recruit a subset of ACBD3 to intracellular membranes.

We thus considered that ACBD3 MWT^374-376^ could be important for interactions with golgin-45 and giantin. We performed GFP-nanobody immunoprecipitation with a negative control GFP, GFP-ACBD3 and GFP-ACBD3-MWT^374-376>AAA^ (Figure 6C). Immunoprecipitates were blotted for endogenous giantin and golgin-45. No interaction was observed with GFP alone, and GFP-ACBD3 interacted with both golgin-45 and giantin. Interestingly, the MWT^374-376>AAA^ mutation that prevents the localisation of ACBD3 to the Golgi apparatus does not interact with either of the golgins. Therefore the second mechanism for Golgi recruitment of ACBD3 is between the MWT^374-376^ residues of ACBD3 and two golgins: golgin-45 and giantin.

### There are two sequential mechanisms for ACBD3 recruitment to the Golgi apparatus

We have demonstrated two mechanisms by which ACBD3 is recruited to the Golgi apparatus. The first is by SEC22B and SCFD1, which participate in a SNARE complex, and the second is by a redundant set of golgins. To experimentally interrogate whether these mechanisms are redundant or sequential, we tested for GFP-ACBD3 binding to the golgins in the context of SCFD1-KO (Figure 7). Interestingly, both golgin-45 and giantin were less efficiently co-immunoprecipitated upon loss of SCFD1. We have therefore demonstrated that recruitment of ACBD3 via SCFD1 is upstream of the interaction with the golgins, and ACBD3 is recruited to the Golgi in a two-step process.

**Figure 7.**
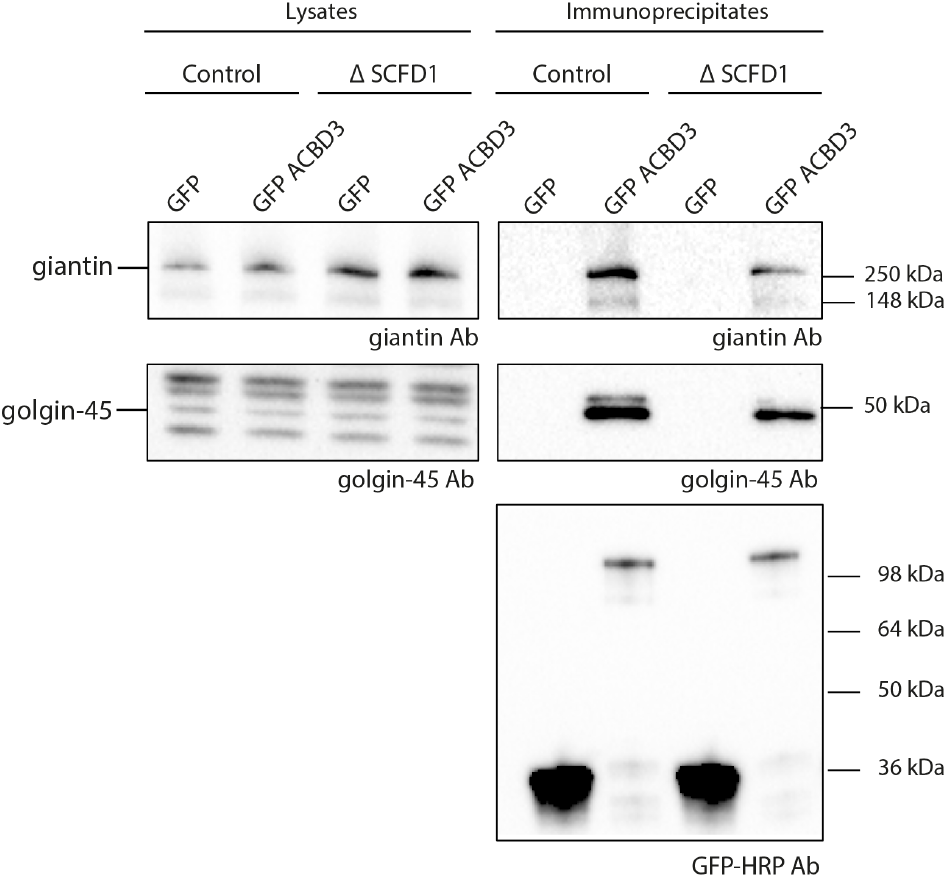
The loss of SCFD1 affects the interaction of ACBD3 with giantin and golgin-45. GFP alone and GFP-ACBD3 were overexpressed in WT or KO SCFD1 HEK-293T cells. Immunoprecipitation experiments were performed and immunoprecipitates were blotted for endogenous giantin and golgin-45. The experiment was performed three times independently.

## Discussion

ACBD3 is a key regulator of the metazoan Golgi apparatus, yet the mechanism of its recruitment to the Golgi was unclear. Using a combination of biochemistry and genetics, we have identified two mechanisms by which ACBD3 is recruited to the Golgi apparatus. The first mechanism is through the SM protein SCFD1, which participates in a SNARE complex with SEC22B. SCFD1 and SEC22B interact with ACBD3 through the GOLD domain on the C-terminus of ACBD3. We also have identified a pair of redundant golgins, giantin and golgin-45, that are important for recruiting ACBD3. We mapped the interaction to a tripeptide MWT^374-376^ sequence in the UR of the GOLD domain of ACBD3. We demonstrate that the interaction with the golgins also depends on the presence of SCFD1, placing the SCFD1-ACBD3 interaction upstream of the golgin-ACBD3 interactions. This demonstrates a tightly regulated process allowing ACBD3 to be recruited to the Golgi with high spatiotemporal regulation.

Combining our data, we postulate a new model for the recruitment of ACBD3 to the Golgi apparatus (Figure 8). Prior to the fusion of a vesicle with the Golgi apparatus, the SNARE-SM complex includes at least four SNAREs and the SM protein SCFD1. The presence of this SNARE-SM complex recruits ACBD3 to the localisation of the incoming vesicle at the *cis*-Golgi. We propose that this ‘activates’ ACBD3, perhaps by mediating a conformational change exposing the MWT^374-376^ motif in the UR. Indeed, ACBD3 has been demonstrated to have multiple different conformations in solution^39,40^. The activated ACBD3 can thus interact with the golgins, retaining the ACBD3 at the *cis*-Golgi beyond the time of the transient *trans*-SNARE complex. The residency time of ACBD3 at the Golgi will therefore be dictated by the affinity to the golgins. Upon recruitment to the Golgi, ACBD3 will co-recruit its effectors, including PI4KIIIβ. These effectors will appropriately modify the incoming vesicle membrane as it arrives in the Golgi apparatus.

**Figure 8.**
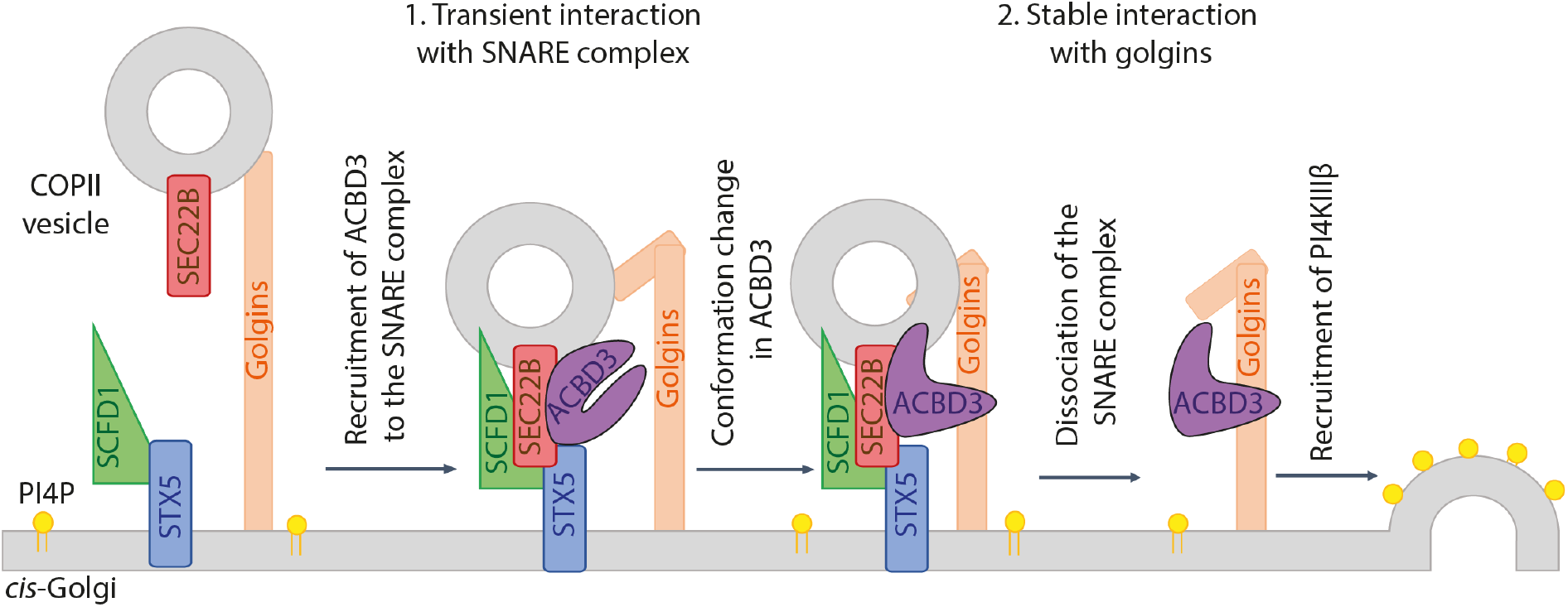
Model: ACBD3 recruitment to the cis-Golgi is achieved through two independent, sequential mechanisms.

One potential caveat with our model is our observation that golgin-45 can recruit ACBD3 to the mitochondrial membrane. However, ACBD3 will exist in both conformations. Our experiment only shows a proportion of total ACBD3 recruited to the mitochondria, and additionally, the mitochondrial golgin-45 is overexpressed. Thus we conclude this mitochondrial-recruited ACBD3 only represents a fraction of the ACBD3.

There have been multiple other models for ACBD3 recruitment to the Golgi apparatus. Based on interaction data, ACBD3 was proposed to be recruited by giantin^32^. Our data and others demonstrate that although we can recapitulate an interaction with giantin, it is not solely responsible for the recruitment of ACBD3 to the Golgi apparatus. ACBD3 has also been proposed to be recruited to the Golgi apparatus by ARL5B^35^. Loss of ARL5B caused 50% of ACBD3 Golgi localisation to be lost. ACBD3 and ARL5B were demonstrated to interact by proximity proteomics. However, immunoprecipitation experiments between ARL5B and ACBD3 did not demonstrate an interaction. Therefore the contribution of ARL5B the recruitment of ACBD3 to the Golgi apparatus seems to be particularly to the *trans*-Golgi, where ARL5B localises. The direct relationship between ARL5B and ACBD3 in the Golgi apparatus remains uncertain; we don’t know if ARL5B recruits ACBD3 to the Golgi directly or if it is upstream. Nonetheless, exploring how this interaction connects with the mechanisms discussed here might reveal an intriguing additional level of control

One factor we have not considered in our experimental approach or model is the oligomerisation of ACBD3. ACBD3 has been demonstrated to oligomerise in the presence of C18:1-CoA or C16:0-CoA^30^. Our data suggest that the binding of ACBD3 to C18:1-CoA, C16:0-CoA or related fatty acyl chains and the subsequent oligomerisation does not affect its recruitment to the Golgi as deletion of the ACBD domain does not prevent Golgi localisation. It is possible that this oligomerisation regulates the conformational change of ACBD3, its effector recruitment, or represents an additional functional role of ACBD3, which could be explored in future studies.

Together, this study showed a highly spatiotemporally-regulated mechanism for the recruitment of ACBD3 to the Golgi apparatus. We discovered previously unidentified interactors and a two-step mechanism of recruitment. Many of the identified putative novel ACBD3 interactors were not important for its recruitment to the Golgi and may represent uncharacterised ACBD3 effectors, which could be investigated in future studies. We also identified a novel MWT^374-376^ motif important for interaction with golgins; further work could investigate if this is specific to ACBD3 or represent a common shared mechanism for protein recruitment to the Golgi.

## Methods

### Antibodies and Other Reagents

The following primary antibodies were used for IF: rabbit anti-ACBD3 (ab134952, 1:200), sheep anti-TGN46 (Bio-rad AHP500G, 1:1000), mouse anti-GM130 (BD 610822, 1:200), mouse anti-PI4KIIIβ (BD Bioscience, 1:500), rat anti-HA (Roche 11867423001, 1:300), mouse anti-EEA1 (BD biosciences 610456, 1:500). Alexa Fluor Dyes (1:1000) were purchased from Invitrogen (Donkey anti-rabbit 488: A32790; Donkey anti-sheep 647: A21448; Donkey anti-mouse 488: A32766; Donkey anti-mouse 647: A32787, Donkey anti-rat 647: A48272).

The following primary antibodies were used for WB in this study: mouse anti-GFP HRP conjugate antibody (Miltenyi Biotec 130-091-833, 1:1000), rabbit anti-golgin-45 (ab155510, 1:1000), rabbit anti-giantin (HPA011008, 1:1000), rabbit anti-SCFD1 (ab86594, 1:1000). Goat HRP-conjugated secondary antibodies (1:5000) were purchased from Abcam (anti-mouse: ab205719; anti-rabbit: ab205718).

The following dyes were obtained from these vendors: Janelia Fluor® HaloTag® Ligand 646 (GA112A; Promega), DAPI (D21490; Invitrogen), HCS CellMask™ Blue Stain (H32720; Invitrogen).

The following antibiotics were used to select cell lines: Puromycin (1 μg/ml for HeLa cells and μg/ml for HEK293T cells; A1113803; Gibco) and Blasticidin (15 μg/ml for HeLa cells and 5 μg/ml for HEK293T cells; A1113903; Gibco).

### Plasmids

pHaloTag-C1 and pHaloTag-N1 were generated by replacing the eGFP in Clontech vectors with HaloTag using Gibson assembly (E2621L; New England Biolabs). HaloTag-ACBD3, SCFD1-HaloTag and HaloTag-SEC22B were cloned by Gibson assembly using a synthetic codon optimised version of the gene of interest (Integrated DNA Technologies), cloned in-frame upstream or downstream of HaloTag in pHaloTag-C1 or pHaloTag-N1. GFP-ACBD3 was generated with the same approach except that peGFP-C1 vector was used.

For the alanine scanning approach in the unique region of ACBD3, alanine mutations were introduced into the HaloTag-ACBD3 vector by using the Q5 Site-Directed Mutagenesis Kit from New England Biolabs. The Q5® Hot Start High-Fidelity 2X Master Mix (M0494S) was used along with custom mutagenic primers to introduce substitutions of target residues in ACBD3 by PCR (Halo-ACBD3 368-370 (VIA^368-370>AAA^), 371-373 (APS^371-373>AAA^), 374-376 (MWT^374-376>AAA^), 377-379 (RPQ^377-379>AAA^), 380-382 (IKD380-382>AAA), 383-385 (FKE383-385>AAA), 386-388 (KIQ386-388>AAA).

The PCR product was used for reaction with the Kinase-Ligase-DpnI (KLD) enzyme mix (M0554S). Truncations of GFP-ACBD3 (1-184, 1-372, 1-400, 328-528, 368-528, 390-528) and HaloTag-SEC22B (isolated longin and SNARE and transmembrane domains) were also generated with the Q5 Site-Directed Mutagenesis Kit.

BirA*-HA-MAO, and golgin-45-HA-MAO were a kind gift from the Sean Munro lab. BirA*-HA-MAO was generated as described previously^50^. Golgin-45-HA-MAO was generated by >90% of the cell population was further confirmed through flow Gibson assembly using the HA-MAO vector digested with NheI/KpnI and golgin-45 cDNA (missing its C-ter, amino acids 1 to 368) purchased from Origene.

Cas9 viral expression backbone was a kind gift from Paul Lehner lab as well as the packaging vectors pMD.G and pCMVR8.91.

pKLV-U6gRNA(BbsI)-PGKblast2ABFP vector was generated by replacing the puromycin resistant gene in pKLV-U6gRNA(BbsI)-PGKpuro2ABFP (Addgene plasmid #50946) with blasticidin via Gibson assembly.

Plasmids and primers used in this work are available upon request. All constructs were sequenced to verify their integrity.

### Cell lines and lentiviral particles production

All cell lines were grown in DMEM high glucose (D6429; Sigma-Aldrich) supplemented with 10% fetal bovine serum (FBS) (F7424; Sigma-Aldrich) and MycoZap™ Plus-CL (VZA-2012; Lonza). They were kept at 37°C in a humidified 5% CO_2_ atmosphere.

HeLa cells were already available in the lab; HEK293T cells were a gift from Janet Deane (CIMR, University of Cambridge, UK); Lenti-X™ 293T cells were obtained from Takara Bio (632180).

Lenti-X™ 293T cells were used to package pKLV-U6gRNA(BbsI)-PGKpuro2ABFP or pKLV-U6gRNA(BbsI)-PGKblast2ABFP vectors encoding guide RNAs into lentiviral particles as previously described^51^. The IDT Alt-R® CRISPR-Cas9 guide RNA tool and CRISPOR (http://crispor.tefor.net/) were used to custom design two guide sequences per gene of interest (except for GOLGB1/giantin and BLZF1/golgin-45 where only one guide sequence was used, see Table S2). Viral supernatants were harvested after 48 h, filtered through a 0.45 μm filter, and when needed, concentrated down 10 or 30 times using the Lenti-X Concentrator (631232; Takara Bio). Supernatants were kept at -80°C prior to being directly applied to target cells which were then spun at 700 xg for 1 h at 37°C. When possible, cells were transiently selected with the appropriate antibiotic 48 h post-transduction.

Stable HeLa Cas9 and HEK293T Cas9 expressing cell lines were generated by infecting cells with lentiviral particles carrying Cas9 plasmid DNA (gift from Paul Lehner) followed by selection for blasticidin antibiotic resistance. Cas9 expression on cytometry analysis, by testing loss of cell surface expression of beta-2 microglobulin, upon transduction with lentiviral particles containing a beta-2 microglobulin targeting guide RNA, using a mouse monoclonal anti-B2M antibody (gift from Paul Lehner).

### GFP trap experiment followed by Mass Spectrometry analysis

1×10^7^ HEK293T cells were seeded onto a 15-cm dish (2 per condition). On day 4, cells were transfected with GFP or GFP-ACBD3 by using PEI (Polyethylenimine branched, 408727, Sigma-Aldrich). The following day, plates were washed once with PBS1X and incubated with 5 ml PBS1X (w/o Ca^2+^ and MgCl_2_) for 10 min at 37°C. Cells were collected and the plates washed 2 times with 5 ml PBS1X. Cells from the same condition were pooled together and were pelleted at 500 xg for 3 min and resuspended with 450 μl of lysis buffer (10 mM Tris pH7.4, 150 mM NaCl, 1 mM MgCl_2_, 0.5% TX100, cOmplete™ EDTA-free Protease Inhibitor Cocktail (Roche, 04693132001). After 30 min incubation on ice, 600 μl of lysis buffer missing TX-100 was added and the lysates were spun for 20 min at 17 000 xg at 4°C. Supernatants were incubated for 1 h with 20 μl prewashed GFP-Trap® Agarose beads (Proteintech, gta-10). Beads were pelleted and washed 2 times with 1 ml of lysis buffer containing 0.2% TX-100 for 5 min on a turning wheel at 4°C followed by 4 quick washes with 1 ml of lysis buffer missing TX-100. 50 μl of Laemmli buffer + DTT (Bio-Rad, 1610747) was added to the beads and samples were boiled for 10 min at 95°C. Beads were removed and the eluates were analysed on a Q Exactive Plus Orbitrap Mass Spectrometer (Thermo Scientific). Protein hits were filtered for Golgi-associated proteins as per Uniprot annotations. This experiment was performed 3 times independently. See table S1 for post-analysis data set.

### Immunoprecipitation experiments

4×10^6^ HEK293T cells were seeded per condition onto a 10-cm dish. On day 3, cells were transfected with GFP alone or GFP-ACBD3 (wild-type, truncations and mutants) together with SCFD1-HaloTag or HaloTag-SEC22B (and isolated domains) by using PEI. Cells were incubated overnight with complete DMEM containing JF646 HaloTag ligand (20 nM; GA112A; Promega). The GFP protein isolation was performed as mentioned above except that only 10 μl of GFP-Trap® Agarose beads were used which were blocked for 1 h with 3% BSA at 4°C. Beads were washed 3 times with 1 ml of lysis buffer containing 0.2% TX-100 (one quick wash and 2 washes of 5 min) followed by one quick wash with 1 ml of PBS1X. 50 μl of Laemmli buffer + DTT was added to the beads and samples were boiled for 10 min at 95°C. 1.25 μl of the lysates (0.125%) and 20-25 μl of the immunoprecipitates were resolved on a gradient Tris-Glycine acrylamide gel. The gel was directly imaged at 647 nm to detect SCFD1-HaloTag or HaloTag-SEC22B (ChemiDoc Imaging System, Bio-Rad). The gel was then transferred to a PVDF membrane which was blocked with 5% skimmed milk and incubated with anti-GFP HRP conjugate antibody overnight at 4°C. Membranes were washed with PBS-T in-between steps, and finally, their immunoreactivity was visualised using Clarity (1705061; Bio-Rad) or WesternBright Sirius (K-12043-D10; Advansta) ECL substrate, in the ChemiDoc Imaging System (Bio-Rad). When bands were quantified, Fiji was used.

For the immunoprecipitation experiments investigating the interaction of ACBD3 with endogenous golgin-45 and giantin, HEK293T cells were transfected with GFP alone, GFP-ACBD3 WT and MWT^374-376>AAA^ and the experiment was carried out as described above except that 60 μl of Laemmli buffer + DTT was added to the beads. 15 μl of the lysates and 20 μl of the immunoprecipitates were resolved on a 8% Tris-Glycine acrylamide gel and transferred to a PVDF membrane. Membranes were incubated with an anti-golgin-45 or anti-giantin antibody overnight at 4°C, followed by a 1 hour incubation with a secondary antibody at room temperature.

For the immunoprecipitation experiments investigating the interaction of ACBD3 with endogenous golgin-45 and giantin in SCFD1 KO cell lines, guide RNAs targeting SCFD1 were cloned into pKLV-U6gRNA(BbsI)-PGKpuro2ABFP and viral particles were produced as described above. For the SCFD1 KO condition, 4 times 2×10^5^ HEK293T Cas9 cells were infected with 250 μl of a 10x concentrated lentiviral supernatant in a 6-well plate. 72 h post-transduction, cells of the same condition were pooled together, replated onto two 10 cm dishes and selected with puromycin. On day 7, cells of each condition were transfected with GFP alone or GFP-ACBD3 WT and the immunoprecipitation experiment was carried out the day after as described above by using the same amount of cells for each condition. 20 μl of the lysates and 20-30 μl of the immunoprecipitates were resolved on a 8% Tris-Glycine acrylamide gel and transferred to a PVDF membrane. Membranes were incubated with an anti-golgin-45 or anti-giantin antibody overnight at 4°C, followed by a 1 hour incubation with a secondary antibody at room temperature.

### Targeted CRISPR knock-out screen

To generate transient KO cells of the different genes identified by mass spectrometry, guide RNAs targeting the gene of interest were cloned into pKLV-U6gRNA(BbsI)-PGKblast2ABFP. Viral particles were produced with Lenti-XTM 293T cells as described above. The HeLa Cas9 stable cell line was then transduced. Briefly, 1×10^4^ HeLa Cas9 cells were infected with 250 μl of lentiviral supernatant in a 96-well plate. 72 h post-transduction cells were replated in a 48-well plate and incubated for an extra 2 days. On day 6, 2×10^4^ of transduced cells were plated onto Matrigel-coated (1:100 in complete DMEM; 354277; Corning) glass coverslips (400-03-19; Academy). On day 8, cells were fixed with 4% PFA for 15 min. Cells were permeabilised with 0.1% TX-100 (T9284; Sigma-Aldrich) and incubated for 1 h with a blocking solution containing 3% BSA (BP9703-100; Sigma-Aldrich). Cells were incubated for 1 h with anti-ACBD3 and anti-GM130 primary antibodies followed by 1 h incubation with appropriate Alexa Fluor Dyes. PBS1X was used to wash cells in-between all steps and DAPI stained (300 nM; D21490 Invitrogen) for 5 min. coverslips were mounted in ProLongTM Gold (P36930; Life Technologies).

### Quantification of ACBD3 localisation at the Golgi apparatus by automated microscopy

6.7×10^3^ HeLa cells were plated in 96-well imaging plates (Viewplates, Perkin Elmer). The next day, cells were transfected using FuGENE® 6 with GFP-ACBD3 WT and the different truncations (1-184, 1-372, 1-400, 328-528, 368-528, 390-528) or with HaloTag-ACBD3 WT and the different alanine mutants. 4 hours post-transfection media was replaced with fresh complete DMEM containing JF646 HaloTag ligand (20 nM; GA112A; Promega) when required. On the following day, cells were fixed with cytoskeletal fixing buffer (300 mM NaCl, 10 mM EDTA, 10 mM Glucose, 10 mM MgCl_2_, 20 mM PIPES pH 6.8, 2% Sucrose, 4% PFA) for 15 min and processed for immunostaining as described above. Anti-GM130 primary antibody was used as well as a HCS CellMask™ Blue Stain (H32720; Invitrogen). Cells were stored in PBS1X at 4°C until imaged.

When using KO cells, 1×10^4^ HeLa Cas9 cells were infected with 25 μl (for SCFD1 and golgin-45) or 45 μl (for giantin) of 10x concentrated lentiviral supernatant in a 96-well plate. 72 h post-transduction, cells were replated onto a 48-well plate and selected with puromycin. On day 6, cells were seeded onto a 96-well Viewplate and, on day 8, immunostaining was carried out by using anti-GM130 and anti-ACBD3 primary antibodies and a HCS CellMask™ Blue Stain.

Images were acquired on a CellInsight CX7 high-content microscope (Thermo Fisher Scientific) using a 20× objective (NA 0.45, Olympus). The Cell Mask was used for automated focussing and cell detection. 40 to 60 fields were imaged in three wells for each condition. Each experiment was performed 3 to 5 times independently. The general intensity measurement algorithm from the HCS Studio software was used as a template with parameters adapted to our experiments. Objects (the Golgi apparatus labelled with GM130 antibody, ACBD3 distribution) were identified using the isodata or the triangle thresholding methods and transfected (GFP-ACBD3 and HaloTag-ACBD3) cells were selected with a fixed area and with a fixed average intensity. Masks of the cell (by using the Cell Mask labelling) and of the Golgi (by using the GM130 labelling) were created with the Circ function. The average intensity of ACBD3 in the total cell and at the Golgi were exported to Excel and the ratio calculated. The intensity of HaloTag or GFP alone was then subtracted to all values. Values from the technical repeats were averaged, and three to five biological repeats were carried out. To account for variance in ACBD3 levels in the KO samples, experiments were normalised to ACBD3 levels at the Golgi.

### Immunofluorescence and Microscopy

For immunofluorescence microscopy, 4×10^4^ HeLa cells were plated onto Matrigel-coated glass coverslips in a 24-well plate. FuGENE® 6 was used to transfect constructs encoding proteins of interest. 4 hours post-transfection media was replaced with fresh complete DMEM containing JF646 HaloTag ligand when required. On the following day, cells were fixed with cytoskeletal fixing buffer and immunostaining was carried out as described above.

For KO of specific genes, 1×10^4^ HeLa Cas9 cells were infected with 250 μl of lentiviral supernatant in a 96-well plate. 72 h post-transduction, cells were replated onto a 48-well plate and selected with puromycin. On day 6, cells were seeded onto Matrigel-coated glass coverslips in a 24-well plate and immunostaining was carried out as described above. Standard epifluorescence images were obtained with an AxioImagerZ2 microscope (Zeiss) equipped with an Orca Flash 4.0v2 camera (Hamamatsu), an HXP 120W light source and a 100x 1.4 NA Plan-Apochromat objective, all under the control of ZEN Blue software (Zeiss).

To analyse the colocalisation of ACBD3 with different compartment markers (TGN46, GM130 and EEA1), cells were imaged on a LSM880 confocal with Airyscan (Zeiss) with a 63x 1.4NA Plan-Apochromat objective. Images were Airyscan processed with a fixed strength parameter set at 6. Colocalisation analysis were performed using the Fiji software plugin Coloc 2 to determine the Pearson’s correlation coefficient among two channels from three independent experiments. The Fiji software plugin Plot profile was used to generate the fluorescence line profiles.

### RNA extraction and qPCR

Total RNA was isolated from cultured HeLa Cas9 WT or KO cells (35mm dish), using the RNeasy Mini Kit (74104; Qiagen) according to the manufacturer’s instructions. RNA quantification was performed using NanoDrop One (Thermo Scientific). Strand cDNA was generated by priming 1 μg of total RNA with a oligo (dT)16/random hexamers mix, using the High-Capacity RNA-to-cDNA™ Kit (4387406; Applied Biosistems), following manufacturer’s instructions. cDNA templates were diluted 50-fold and 5 μL was used with specific oligos spanning 2 continuous exons (Table S3) along with the PowerUpTM SYBRTM Green Master Mix (A25741; Applied Biosistems) for the qPCR reaction. All reactions were performed in 3 technical replicates using the CFX96 Touch Real-Time PCR Detection System (Bio-Rad). Data was analysed according to the 2^-ΔΔCT^ method^52^.

### Statistical Analysis

Statistical analysis was performed using Python 3.7 and statistical significance was considered when P<0.05. Comparisons were made using Student’s t-test or Multiple Comparison of Means - Tukey HSD, FWER=0.05. All quantitative data are expressed as mean ± standard deviation of at least three independent experiments.

## Acknowledgements

We thank all lab members for their help and useful comments. We thank Sean Munro for advice, support and reagents. We thank Paul Lehner and Dick van den Boomen for molecular biology reagents and advice.

This research was supported by the CIMR Flow Cytometry Core Facility, the CIMR Microscopy Facility and CIMR Mass spectrometry facility. In particular, we wish to thank Matthew Gratian, Reiner Schulte, Jack Houghton, Robin Antrobus and Gabriela Grondys-Kotarba for their advice and support in imaging, proteomics, bioinformatics, flow cytometry and cell sorting. D.G. and D.S. are funded by a Sir Henry Dale Fellowship awarded to D.G. from the Wellcome Trust/Royal Society (Grant 210481).

For the purpose of open access, the author has applied a Creative Commons Attribution (CC BY) licence to any Author Accepted Manuscript version arising.

We thank Steve Royle and the Royle Lab for the aviliblity of the LAT_E_X template used for typesetting.

We thank Gillian Griffiths, Margaret Robinson, Daniel Fazakerley, Laura Pellegrini and the Gershlick Lab for reading the manuscript, helpful discussions and advice.

## Supplementary Information

**Figure S1.**
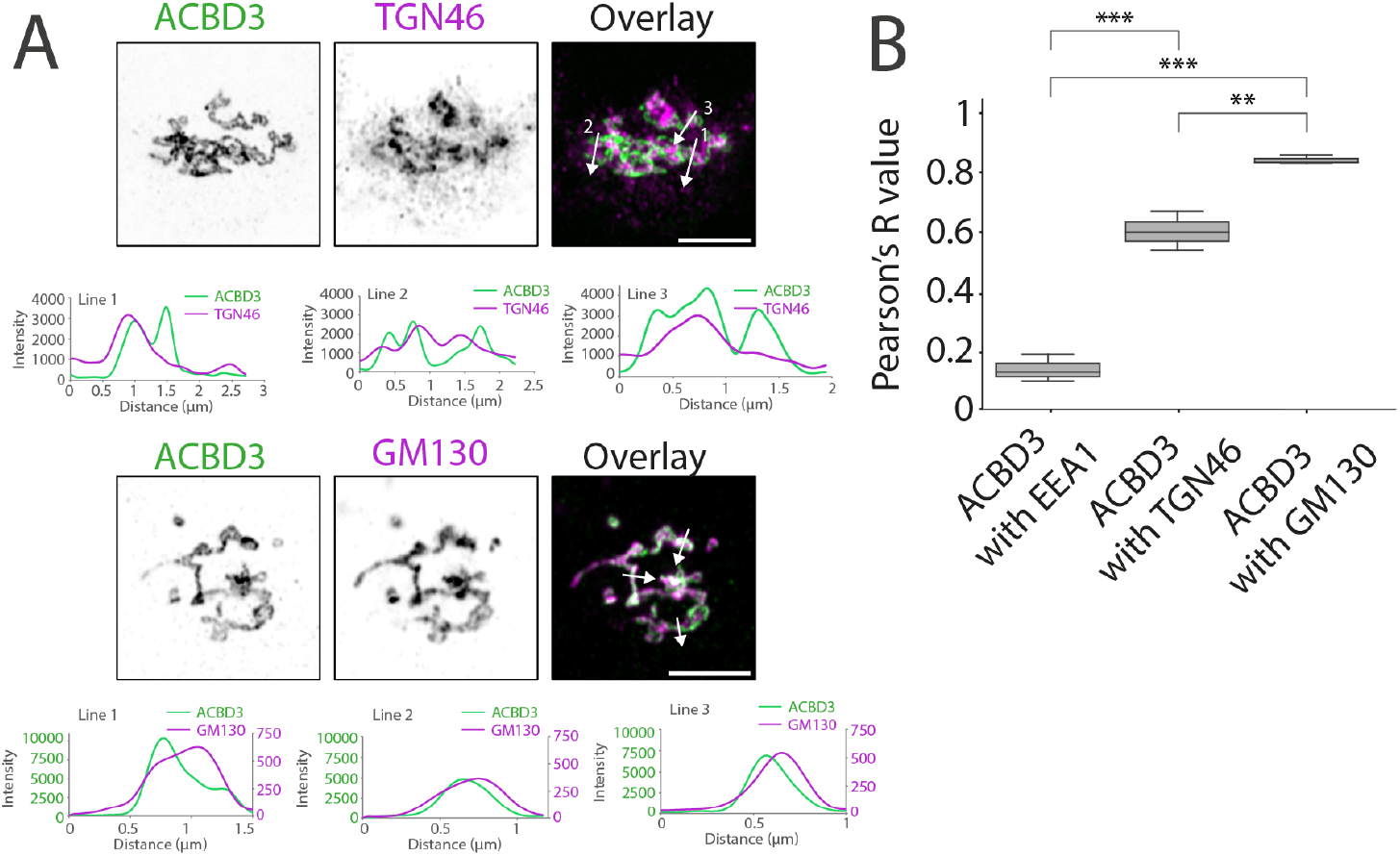
ACBD3 localises to the Golgi apparatus. (**A**) Airyscan imaging of HeLa cells colabelled with ACBD3 and TGN46 (*trans*-Golgi marker) or GM130 (*cis*-Golgi marker). Scale bar: 5 μm. Fluorescence line profiles show the enrichment of ACBD3 at both the *cis* and *trans*-Golgi. (**B**) Quantification of A. The Pearson’s correlation coefficient was calculated from 3 independent experiments (number of cells per experiment). EEA1, an endosomal marker, was used as a negative control. Tukey’s multiple comparisons test (HSD, FWER=0.05) was performed. **P≤0.01; ***P≤0.001.

**Figure S2.**
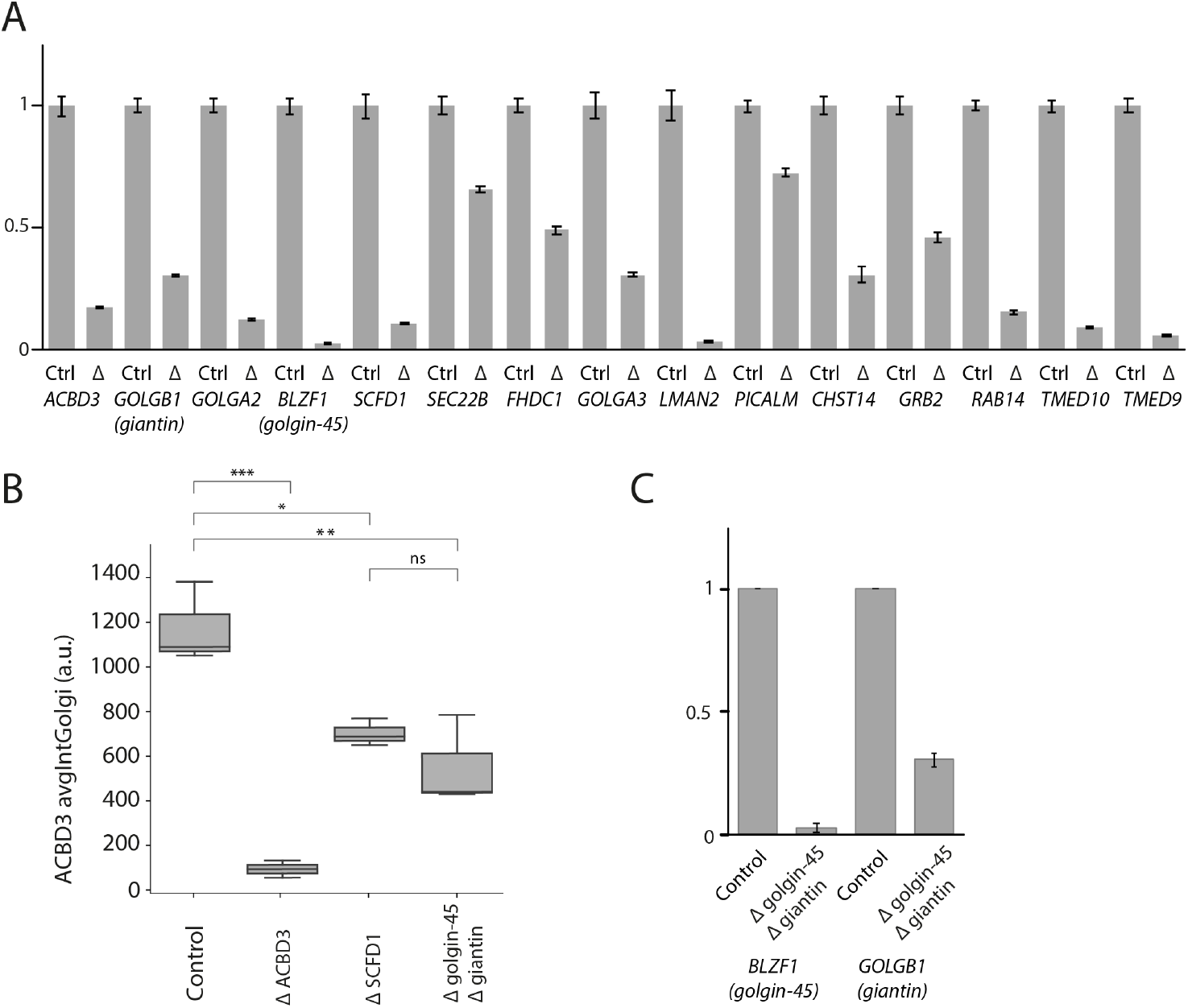
Validation of guide RNA KO efficiency by qRT-PCR and quantification of ACBD3 localisation at the Golgi apparatus in different KO cell lines. (**A**) Validation of guide RNA KO efficiency by qRT-PCR. Data from all conditions was internally normalised to GAPDH and TBP expression and is represented as fold change of control HeLa Cas9 cells. Error bar = SD of 3 technical repeats. (**B**) Quantitative analysis with a CellInsight CX7 high-content microscope of the Golgi localisation of ACBD3 in HeLa cells knocked-out for ACBD3, SCFD1 or double knocked-out for the two golgin, giantin and golgin-45. The average intensity of ACBD3 at the Golgi is indicated (a.u). The experiment was performed 3 times independently. Tukey’s multiple comparisons test (HSD, FWER=0.05) was performed. *P≤0.05; **P≤0.01; ***P≤0.001. ns: not significant. (**C**) Validation of the double KO of the two golgins, giantin and golgin-45 in HeLa Cas9 cells. Data was internally normalised to GAPDH and TBP expression and is represented as fold change of control HeLa Cas9 cells. Error bar = SD of 3 independent experiments.

